# Exploring the emergence of morphological asymmetries around the brain’s Sylvian fissure: a longitudinal study of shape variability in preterm infants

**DOI:** 10.1101/2022.07.15.500199

**Authors:** H de Vareilles, D Rivière, M Pascucci, Z Sun, C Fischer, F Leroy, ML Tataranno, MJNL Benders, J Dubois, JF Mangin

## Abstract

Brain folding patterns vary within the human species, but some folding properties are common across individuals, including the Sylvian fissure’s inter-hemispheric asymmetry. Contrarily to the other brain folds (sulci), the Sylvian fissure develops through the process of opercularization, with the frontal, parietal and temporal lobes growing over the insular lobe. Its asymmetry may be related to the leftward functional lateralization for language processing, but the time-course of these asymmetries’ development is still poorly understood. In this study, we investigated refined shape features of the Sylvian fissure and their longitudinal development in 71 infants born extremely preterm (mean gestational age at birth: 26.5 weeks) and imaged once before and once at term-equivalent age (TEA). We additionally assessed asymmetrical sulcal patterns at TEA in the perisylvian and inferior frontal regions, neighbor to the Sylvian fissure. While reproducing renown strong asymmetries in the Sylvian fissure, we captured an early encoding of its main asymmetrical shape features, and we observed global asymmetrical shape features representative of a more pronounced opercularization in the left-hemisphere, contrasting with the previously reported right-hemisphere advance in sulcation around birth. This added novel insights about the processes governing early-life brain folding mechanisms, potentially linked to the development of language-related capacities.

**Highlights:**

- Shape features can be isolated to describe quantitatively the development of the Sylvian fissure
- Strong asymmetries are encoded as soon as 30 weeks of post-menstrual age
- The process of opercularization is more pronounced in the left hemisphere

## Introduction

Compared to other species, one of the specificities of the human brain lies in its anatomical asymmetries, implying a different macroscopic configuration of the left and right hemispheres. Not all asymmetries which have been reported in the human are human-specific (Amiez et al., 2021), yet a consensus is reached on some of them, including the Yakovlevian torque (Hou et al., 2019; frontal forward warping of the right hemisphere and dorsal backwards warping of the left hemisphere), asymmetries in the Sylvian fissure’s shape (Hou et al., 2019), and a depth asymmetry in the superior temporal sulcus (STS) (Leroy et al., 2015).

These human-specific asymmetries have been described through the study of cortical folds (including fissures and sulci) since these objects can be reliably identified for comparisons within and between individuals. In particular, in the adult brain, the Sylvian fissure has been reported to be shorter and more anteriorly bending in the right hemisphere than the left one (Lyttleton et al., 2009), and right-hemisphere STS have been reported to be deeper than left ones in their middle part, under Heschl’s gyrus (Leroy et al., 2015), with specific relationships to genetic constraints (Le Guen et al., 2019). These asymmetries are considered to be linked to the functional lateralization of brain networks.

Indeed, the most renown functional lateralization of the human brain is the one observed for language processing, which can be roughly dissociated between a ventral stream for speech perception and a dorsal stream for speech production (Hickok and Poeppel, 2007), both involving the perisylvian regions (brain regions located around the Sylvian fissure). The primary auditory cortex is located in Heschl’s gyrus, within the temporal wall of the Sylvian fissure. Just behind, the *planum temporale* ensures the spectrotemporal analysis of sounds. Posterior to the *planum temporale* is Wernicke’s area, responsible for speech comprehension. Also involved in speech production are Broca’s area, located in the inferior frontal gyrus of the left hemisphere, and the parieto-temporal region, just along the posterior extremity of the Sylvian fissure, which ensures the transformation of sensory encoding of sounds for the motor system. The language ventral stream has been reported to show mostly bilateral activations across hemispheres, while the dorsal stream presents a strong leftward functional asymmetry (Hickok and Poeppel, 2007): about 90% of the general population processes word generation through the left hemisphere (Knecht et al., 2000). The relationship between this strong functional lateralization and the anatomical asymmetries of the perisylvian structures are still under investigation (Sprung-Much et al., 2021).

In parallel, correlations between functional specialization and anatomical properties have already been demonstrated using sulcal pattern as an anatomical landmark (Mangin et al., 2015; Jiang et al., 2021), with recent studies showing interesting population-wise tendencies. For example, an asymmetrical configuration of either the anterior cingulate cortex (inducing a more prominent paracingulate sulcus in the left hemisphere than in the right), or the inferior frontal sulcus’s pattern (with one continuous and one interrupted, interdependently of the hemisphere) seem to be related to a better inhibitory control in children and adults (Tissier et al., 2019). The central sulcus has been reported to show shape asymmetries related to handedness including height considerations about the “hand-knob” (a specific bump of the central sulcus; Sun et al., 2012), and the height of this “hand-knob” seems intertwined with the height of the hand functional activation (Sun et al., 2016). This same study demonstrated anatomo-functional links between sulcal patterns around the Sylvian fissure and a task related to language: the location and size of functional activations induced by silent reading are linked to the pattern of sulci in the precentral gyrus, and along the superior temporal sulcus, depending on the pattern of the neighboring sulci. This leads us to consider that shape asymmetry of folds around the Sylvian fissure might carry out valuable links to function and should be investigated further, including by getting a finer understanding of its early-life development.

This leads us, in a first time, to clarify the developmental phenomenon leading to the formation of the Sylvian fissure and its neighboring sulci. The Sylvian fissure forms through the process of opercularization (Fig. 1): differential growth of the frontal, parietal and temporal cortices and underlying white matter, compared to the insular cortex, which induces the growth of these regions over the insula (Kostović et al., 2019). On the reverse, sulci surrounding the Sylvian fissure form through the classical sulcation process, supposed to rely on several mechanisms among which the larger expansion of cortical grey matter compared to white matter (Llinares-Benadero et al., 2019). In terms of relative chronology, opercularization starts earlier than sulcation, as the start of invagination of the insula happens as soon as 14 to 16 weeks of gestational age (w GA), while the primary sulci first emerge around 22w GA in fetuses (Habas et al., 2012). In terms of anatomical asymmetries throughout development, the right hemisphere shows an advance in sulcation, with an advance of around 2 weeks in the right hemisphere compared to the left (Dubois et al., 2008, Habas et al., 2012).

**Figure 1.**
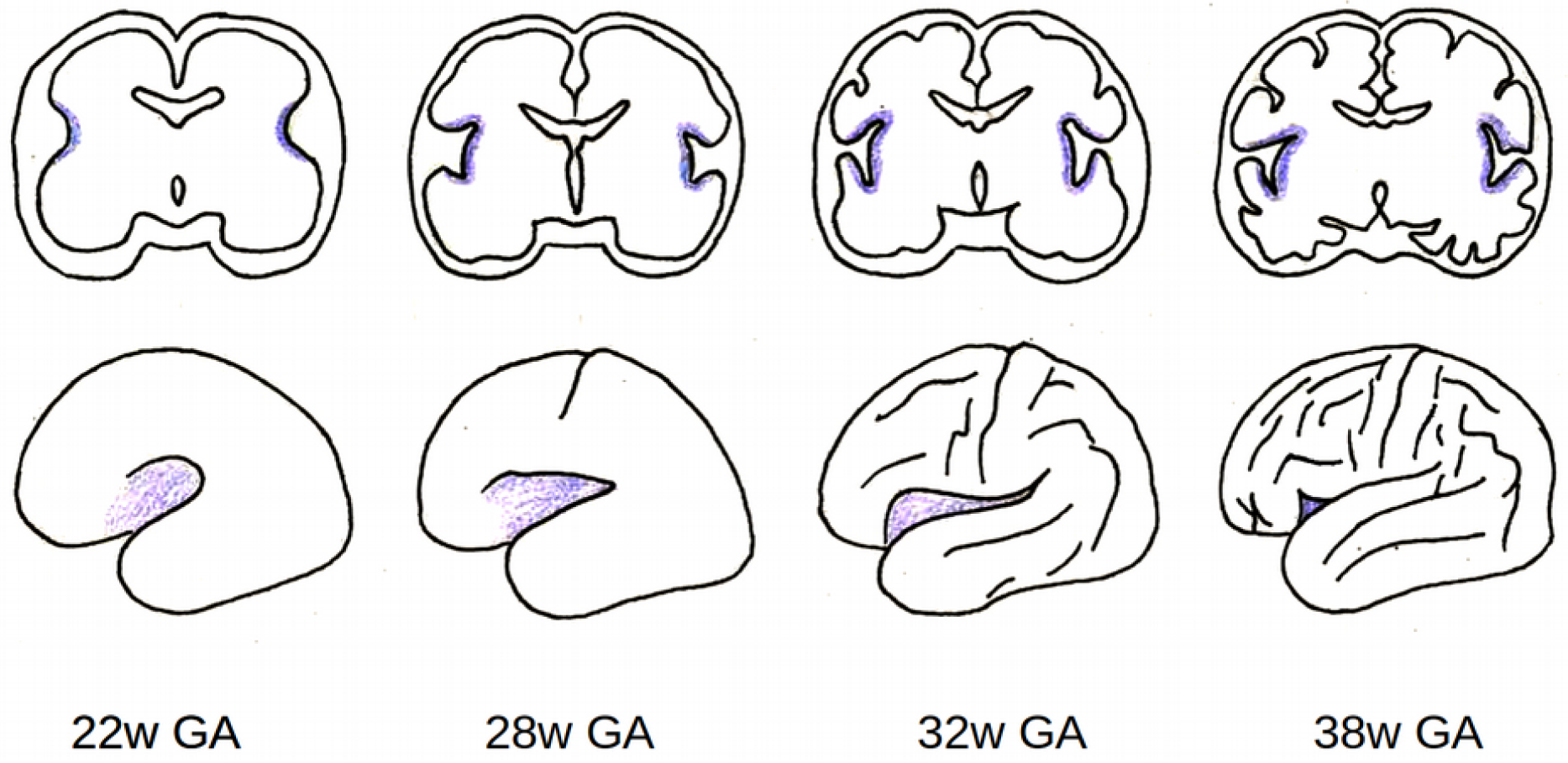
Illustration of the opercularization process between 22 and 38 weeks of gestational age (w GA). Top row: representation of a coronal slice at the level of the insula. Bottom row: volumetric 3D representation, side view of the left hemisphere. Purple: opercularization zone.

Going back to language-related perisylvian regions, the Sylvian fissure’s asymmetries reported in the adult have been observed as being already present in childhood from 7 years of age and adolescence, and to increase with age (Sowell et al., 2002). Asymmetries both in the Sylvian fissure and the STS have been accounted for in the infant brain (Hill et al., 2010; Glasel et al., 2011), and to a certain extent in the preterm (Dubois et al., 2008; Dubois et al., 2010; Kersbergen et al., 2016) and fetal brain (Habas et al., 2012). This raises questions on how these asymmetries emerge in the very young brain. Studying folding mechanisms in the preterm is then particularly interesting as it allows for longitudinal analyses including developmental stages happening before normal-term birth without being impacted by brain modifications induced by birthing (Lefèvre et al., 2016). It should still be noted that prematurity has been linked to altered gyrification (Bouyssi-Kobar et al., 2016; Dubois et al., 2019).

In this context, the purpose of this study was to investigate developmental asymmetries of the Sylvian fissure and regions around it, through MRI acquisitions of a cohort of infants born extremely premature and imaged longitudinally at two ages, once before full-term birth and once at term-equivalent age (TEA). To explore regions that anatomically match the aforementioned language-related regions, we decided to focus on three folding pattern proxies: the Sylvian fissure, the perisylvian region, and the inferior frontal region. The Sylvian fissure and perisylvian region proxies (defined as the congregation of sulci from the inferior frontal, inferior parietal and superior temporal lobes neighboring the Sylvian fissure, belonging to a series of different functional regions) were included to investigate the process of opercularization which modulates the whole folding patterns surrounding the Sylvian fissure. Additionally, we considered the inferior frontal region (selecting sulci from the inferior frontal gyrus) because of its anatomical correspondence to Broca’s region in the left hemisphere and thus its specific relevance in language lateralization.

## Materials and methods

### Cohort

The present study was conducted on a longitudinal retrospective cohort of 71 infants born extremely preterm (gestational age at birth: 26.5 ± 1.0 weeks, range: 24.4 – 27.9 weeks) admitted in the intensive care unit of the Wilhelmina Children’s Hospital in Utrecht (the Netherlands) between June 2008 and March 2013 (Kersbergen et al., 2016). Clinical specificities about this cohort are detailed in Table 1. Approval for this study was granted by the medical ethics committee.

**Table 1.**
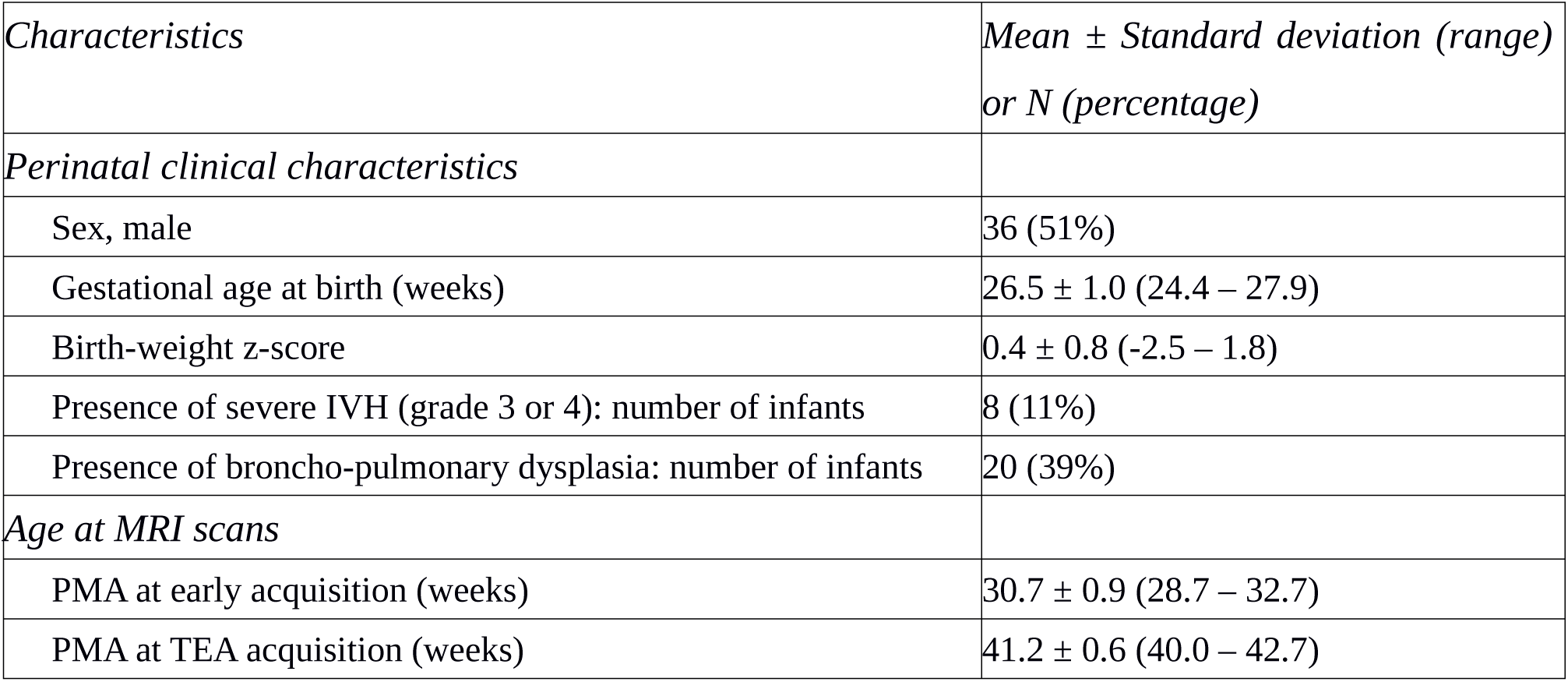
Perinatal clinical characteristics of the study participants

| *Characteristics* | *Mean ± Standard deviation (range) or N (percentage)* |
| --- | --- |
| *Perinatal clinical characteristics* |  |
| Sex, male | 36 (51%) |
| Gestational age at birth (weeks) | $26.5 \pm 1.0$ (24.4 – 27.9) |
| Birth-weight z-score | $0.4 \pm 0.8$ (-2.5 – 1.8) |
| Presence of severe IVH (grade 3 or 4): number of infants | 8 (11%) |
| Presence of broncho-pulmonary dysplasia: number of infants | 20 (39%) |
| *Age at MRI scans* |  |
| PMA at early acquisition (weeks) | $30.7 \pm 0.9$ (28.7 – 32.7) |
| PMA at TEA acquisition (weeks) | $41.2 \pm 0.6$ (40.0 – 42.7) |

### MRI acquisitions

Each subject underwent two MRI acquisitions, one around 30 weeks of post-menstrual age (w PMA) (30.7 ± 0.9, range: 28.7 - 32.7) and one around 40w PMA (41.2 ± 0.6, range: 40.0 - 42.7) corresponding to TEA. The images were acquired on a 3-Tesla MR system (Achieva, Philips Medical Systems, Best, The Netherlands). The protocol included T2-weighted imaging with a turbo-spin echo sequence in the coronal plane (MRI at ∼30w PMA: repetition time (TR) 10,085 ms; echo time (TE) 120 ms; slice thickness 2 mm, in-plane spatial resolution 0.35 × 0.35 mm; MRI at ∼40w PMA: TR 4847 ms; TE 150 ms; slice thickness 1.2 mm, in-plane spatial resolution 0.35 × 0.35 mm).

### Image preprocessing

The MRI brain images underwent preprocessing in order to reconstruct the 3D volumes of the brain as described in (Kersbergen et al., 2016, de Vareilles et al., 2022). The T2-weighted images were first of all segmented in three classes: cerebrospinal fluid, unmyelinated white matter and grey matter using supervised voxel classification (Moeskops et al., 2015). Using the BrainVISA software (https://brainvisa.info) and combining the Morphologist (publicly available) and BabySeg (internal toolbox adapted for infant brain segmentation) toolboxes, we were able to reconstruct the inner cortical surface of both hemispheres and extract items depicting the sulci. A summary of the segmentation pipeline for sulcal extraction is presented Figure 2.A.

**Figure 2.**
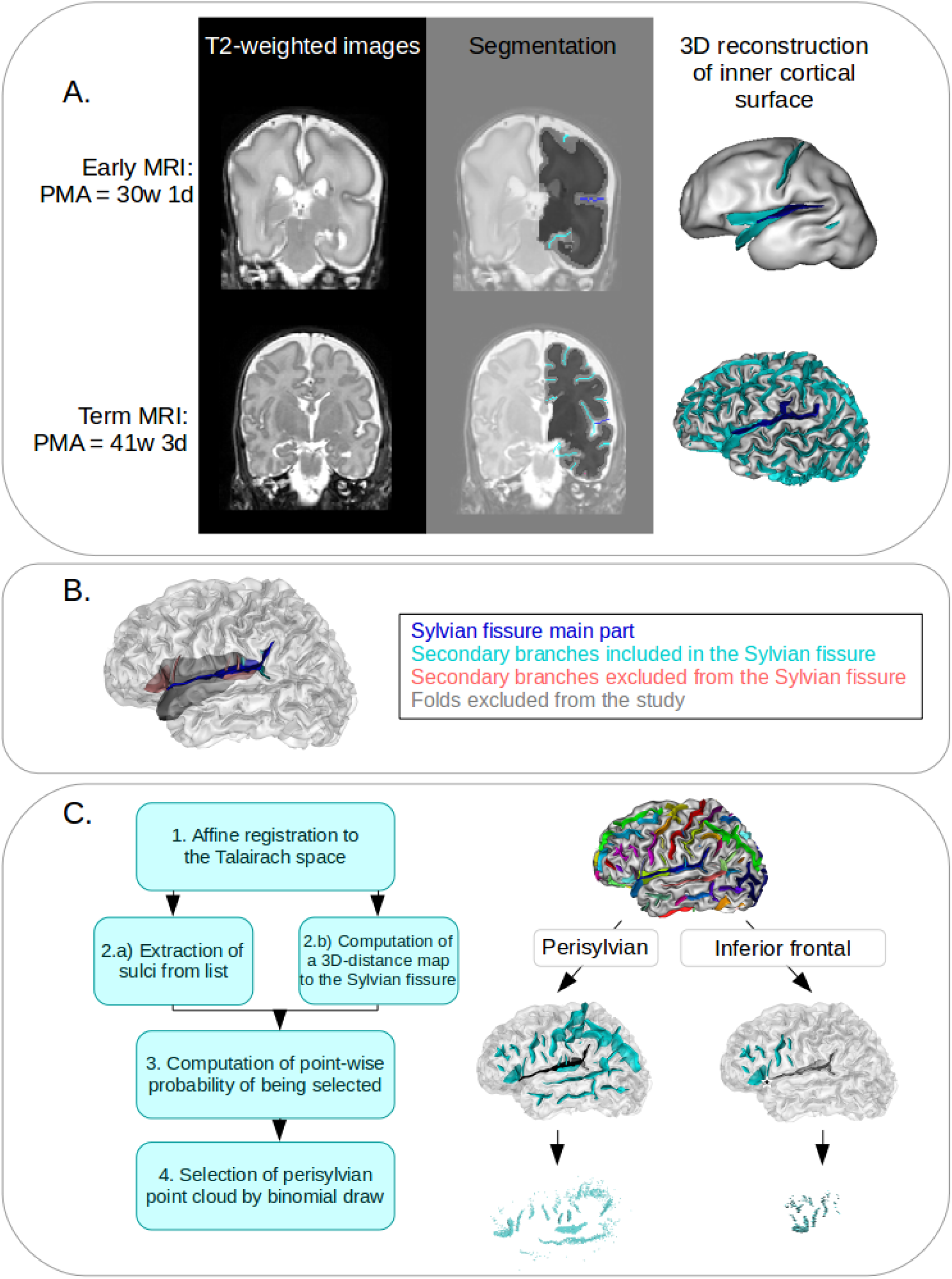
Definition of the study proxies. A. Illustration of the segmentation pipeline. Left: Sample T2-weighted coronal MRI. Middle: Corresponding segmentation on the left hemisphere, showing the boundary between white (dark grey) and grey (light grey) matter. The sulci are represented in cyan, except for the Sylvian fissure, in deep blue. Right: Resulting reconstruction of inner cortical surface. Deep blue ribbon: Sylvian fissure (main branches), cyan ribbons: other sulci. B. Definition of the sulcal elements included and excluded from the Sylvian fissure proxy. Deep blue: main part, cyan: included secondary branches, pink: excluded secondary branches, grey: folds excluded from the study. C. Description of the method for selection of perisylvian and inferior frontal gyrus proxies. Left: Summary of steps to generate the point cloud of a subject on a given hemisphere. Right: Illustration on an infant’s left hemisphere: Top: Full hemisphere with labeled sulci, Middle: Sulci retained based on inclusion list. Cyan: sulci included, black (sulcal element or star): element of the Sylvian fissure used to create the distance map, white: elements of the Sylvian fissure ignored for the distance map computation, Bottom: Resulting point cloud and probabilistic selection of points based on their distance to the relevant Sylvian fissure element.

### Sulcal extraction and labelling

Based on the white matter segmentation and the resulting reconstruction of the inner cortical surface, sulci were materialized as the surface created by the set of voxels equidistant to the walls of two adjacent gyri (Mangin et al., 1995). Once extracted, the sulci were automatically labeled using a Bayesian pattern recognition strategy (Perrot et al., 2011). Finally, the labelling was manually checked and corrected when necessary by one of the authors (HV) using open access software Anatomist (Rivière et al., 2022). In particular and as detailed below, we focused on the labelling of the 1) Sylvian fissure and its branches, 2) sulci of the perisylvian region, and 3) sulci of the infero-posterior region of the frontal lobe.

#### 1) Selection of anatomical landmarks for the Sylvian fissure

In order to study the development of the Sylvian fissure longitudinally, our aim in the selection of its anatomical landmarks was to capture comparable proxies at 30 and 40w PMA. For this reason, we excluded the anterior part of the Sylvian fissure, which was not reliably extracted from 30w PMA brain reconstructions. Then, we selected the Sylvian fissure’s landmarks to include: 1) the main part of the fissure, ascending from the insula, which was given a specific label, 2) the branches stemming from this main part that appear posteriorly to the central sulcus, under the condition that they are anchored at mid-depth or deeper of the main path of the fissure and that they reach the surface of the brain to create a visible sulcus, and 3) ramifications of the main part of the fissure appearing posteriorly to the postcentral gyrus, either on the parietal or temporal lobe. The other branches of the Sylvian fissure which do not meet the criteria (such as the anterior branches diving in the inferior frontal gyrus, the sulci buried in the insula, and the buried sulcus in the *planum temporale*) were labeled *per se* and considered independently from the Sylvian fissure. These criteria were chosen to capture analogous proxies for the Sylvian fissure at 30 and 40w PMA since the excluded sulci were absent from 30w PMA images, while early post-central ramifications were occasionally captured. Figure 2.B illustrates our choice of Sylvian fissure proxy definition in preterm infants at 40w PMA.

#### 2) Selection of anatomical landmarks of the perisylvian region

In order to study the sulcal specificities around the Sylvian fissure, we first considered selecting all sulci neighboring the Sylvian fissure, but this approach was problematic because a number of sulci neighboring the Sylvian fissure reach farther than the perisylvian region (such as the precentral, central and post-central sulci for example). Moreover, the sulcal pattern variability sometimes led sulci which do not usually neighbor the Sylvian fissure to extend up to the perisylvian region. We therefore chose to include sulcal items based on their distance to the Sylvian fissure, by opting for an inclusion of probabilistically pruned point clouds of neighboring sulci, with the pruning weighted by the distance to the Sylvian fissure. First of all, to extract sulcal elements defining the perisylvian region, we defined a list of labels which can be assigned to any sulcus susceptible to neighbor the Sylvian fissure (e.g. inferior frontal sulcus, intraparietal sulcus, superior temporal sulcus; see the complete list of sulci labels in Annex 1). The subsequent steps were carried out independently for each individual and each hemisphere, after affinely registering the brains to the Talairach space.

First, we extracted the sulci corresponding to the predetermined list as a point cloud. We computed a 3-D distance map capturing the Euclidean distance to the Sylvian fissure’s main part (even though we had excluded the Sylvian fissure from the point cloud of selected sulci). We used a binomial draw on each of the cloud points, using a Gaussian probability *p* based on its distance to the Sylvian fissure’s main part. Let X be a point from the point cloud, p(X) its probability of being selected and d(X) its distance to the Sylvian fissure. We defined p(X) as:

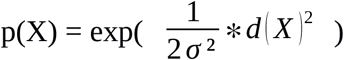

with σ² the variance (i.e. σ the standard deviation), an adjustable constant to constraint the size of the region selected. The σ² parameter was adjusted visually, after examination of the point cloud resulting of a range of different values on two given subjects, with considerations about the ratio of relevant versus irrelevant sulci captured, and validated by visual examination of the resulting point clouds of the whole cohort.

In the perisylvian study, it resulted in σ²=80mm^2^ (σ=8.94mm).

This resulted in a point cloud sampling the sulci neighboring the Sylvian fissure based on their distance to its main part. These steps are summarized on Figure 2.C. This pipeline was only applied to 40w PMA brain reconstructions because of insufficient sulcation at 30w PMA.

#### 3) Selection of anatomical landmarks of the inferior frontal gyrus

The method to focus more specifically on the inferior frontal gyrus was similar to that used for the perisylvian region, but was operated separately and differed on two aspects. First, the initial sulcus list was centered around the area, therefore including the anterior branches of the Sylvian fissure (horizontal ramus, ascending ramus, diagonal sulcus), the inferior frontal sulcus and the inferior and intermediary part of the precentral sulcus. These are classically considered as the anatomical landmarks of Broca’s area, delineating the pars orbitalis, the pars triangularis, and the pars opercularis. Secondly, the sampling of the point cloud was based on the most anterior point of the posterior Sylvian fissure, as it is the junction of its anterior branches in most classical configurations (this point was automatically within the points of the Sylvian fissure based on their spatial coordinates), rather than the whole main part. Once again, we selected σ² by hand after visual examination, resulting in σ²=240mm² (σ=15.49mm). These steps are illustrated on Figure 2.C. Once again, this pipeline was only applied to 40w PMA brain reconstructions because of insufficient sulcation at 30w PMA.

### Pipeline for shape characterization

For each sulcal proxy of this study (even the complex ones relying on point clouds), we used a similar methodology for shape characterization as the one used in a previous study of infants’ central sulci (de Vareilles et al., 2021), derived from a method previously applied to adults (Sun et al., 2016), and based on publicly available pipeline (https://github.com/neurospin/point-cloud-pattern-mining). As detailed below, this included the following steps: folding pattern proxies registration, computation of shape dissimilarity matrix and dimensionality reduction.

First, every subjects’ brain was affinely aligned into the Talairach space, and after extracting the relevant folding pattern proxies from both hemispheres (referred to as proxies in the following), the right ones were mirror-flipped to be comparable to the left ones. In order to capture the shape variability from the whole cohort, a pairwise coregistration using the Iterative Closest Point (Besl & McKay, 1992) was applied to every pair of proxies.

For the perisylvian and inferior frontal regions, the retained proxies presented too much shape variability for simple pairwise registration to ensure anatomical consistency: without further anatomical anchoring, proxies extracted from very dissimilar brains might show optimal registration by matching together incompatible sulci. Therefore, we chose to add items derived from the main part of the Sylvian fissure exclusively for the registration step. For that matter, in the analysis of perisylvian region, we added the whole Sylvian fissure in the point clouds for the registration steps. In the analysis of inferior frontal gyrus, we included only an anterior subpart of the Sylvian fissure (selected with the same method as the original point cloud, but with σ²=480mm²), in order to ensure that this sulcal element would be consistently registered between subjects (including the whole Sylvian fissure would have driven the registration based on its large posterior part, while we were interested in matching the anterior parts). As a result, the registrations were computed with point clouds obtained by including at least a part of the Sylvian fissure in addition to the sulci list before sampling, and the corresponding transformations were then applied to the original point clouds, which did not include the Sylvian fissure (see Sup. Fig S1).

After this step, anatomically consistent pairwise registrations of the initial proxies were achieved, which allowed us to compute the residual Wasserstein distances after coregistration and to store them in a matrix. The resulting matrix therefore captured the shape dissimilarity of the cohort but was of high dimension N_items_×N_items_ with N_items_ the number of proxies considered (N_items_= 71 subjects × 2 hemispheres × number of acquisitions). As the Sylvian fissure folds very early on during brain development, its shape was characterized longitudinally, including the proxies from both 30 and 40w PMA acquisitions (N_items_ = 71 subjects × 2 hemispheres × 2 acquisitions = 284). On the contrary, folding patterns of the perisylvian region and inferior frontal gyrus are almost not visible at 30w PMA, so the shape of the corresponding proxies was investigated exclusively at 40w PMA (N_items_= 71 subjects × 2 hemispheres = 142).

The Isomap algorithm (Tenenbaum et al., 2000) was then applied for dimension reduction, resulting in N_dim_ dimensions capturing the main shape variability from this matrix. This pipeline is summarized in Figure 3. The relevant number of dimensions *d* as well as the number of nearest neighbors *k* retained for the Isomap algorithm were determined according to the methodology developed for the central sulcus in infants (see de Vareilles et al., 2021, Annex 1). The resulting parameters were: *d* = 6 and *k* = 7 for the Sylvian fissure; *d* = 10 and *k* = 7 for the perisylvian region; *d* = 8 and *k* = 7 for the inferior frontal gyrus.

**Figure 3.**
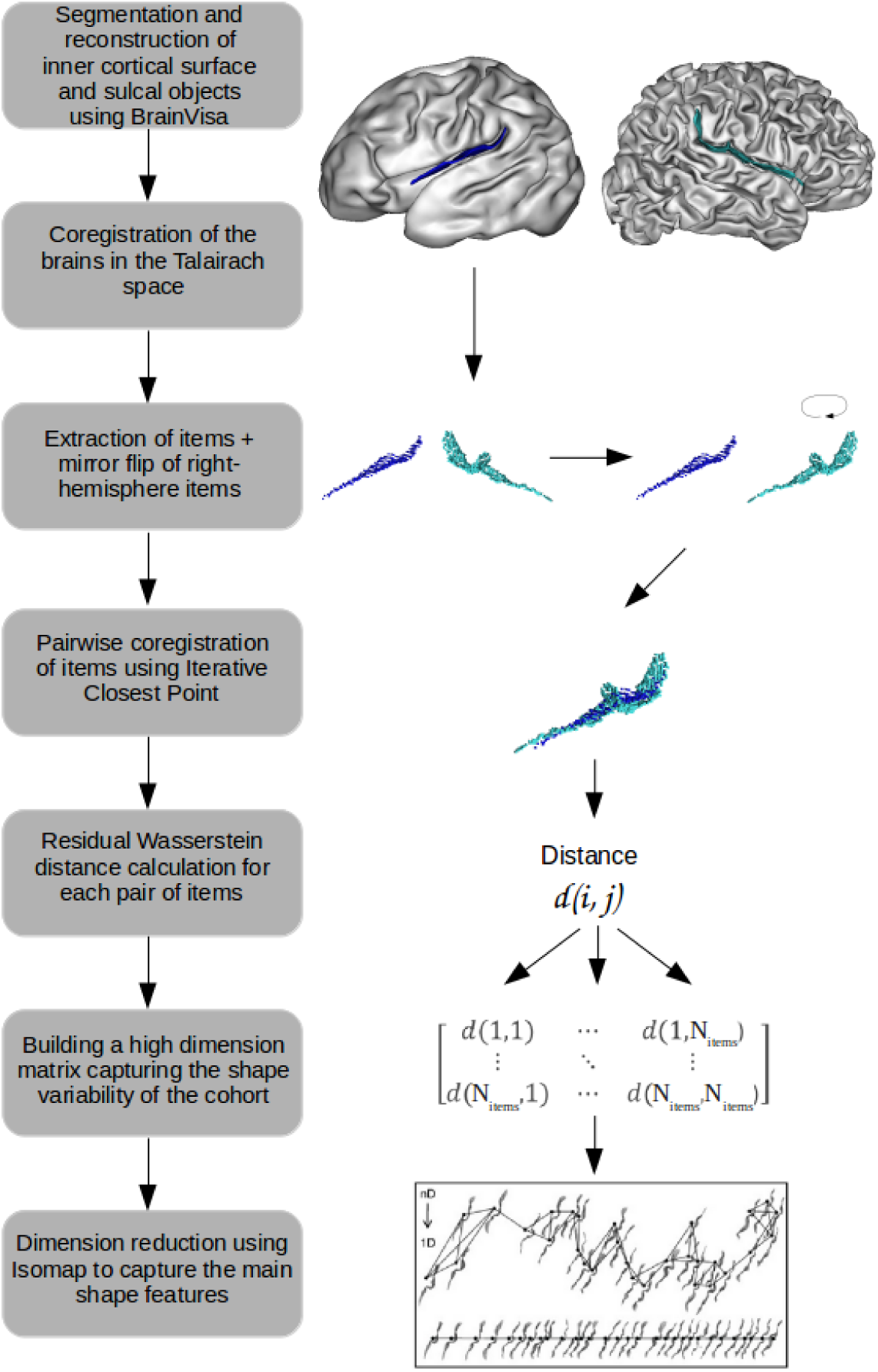
Summary of the general pipeline for shape characterization, illustrated on a left hemisphere 30w PMA Sylvian fissure proxy (deep blue) and a right hemisphere 40w PMA Sylvian fissure proxy (cyan).

### Correction for inter-individual variability in age at MRI

Even though the subjects were aimed to be imaged at the same age (either around 30w or 40w PMA), there was some age differences between the subjects (up to 4 weeks), which might have an impact on the folding shape given the considered periods of development. Therefore, before any additional analysis, we tested if some of the shape variability we captured was linked to the infant’s age by computing the Pearson correlations between positions of each folding pattern proxy on the isomap dimensions and the infants’ age at MRI, combining left and right folding pattern proxies, as presented in Sup. Table S1 (for the Sylvian fissure studied at both 30w and 40w PMA, this test was conducted separately for each age group).

To prevent from considering effects which could be due to this age difference as intrinsic shape variability, we further corrected all the positions on isomap dimensions for age at MRI by considering the residual positions *r*_*n*_ as the shape characteristics of interest in the following numerical analyses (referred to as Isomap positions corrected for PMA). Such corrected positions *r*_*n*_ were computed by solving the following linear model, for the 30 and 40w PMA groups independently:

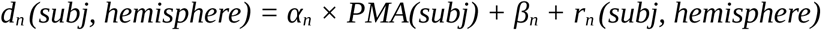

with *d*_*n*_ the raw position of the folding pattern proxies on the n^th^ dimension of the Isomap, *PMA* the post-menstrual age at MRI, *α*_*n*_ and *β*_*n*_ constants estimated by the model. We did not apply this correction to the visualization results, since they were intended to enable us to decipher the overall shape variability captured by the isomap algorithm (even if some of that variability was relayed by age differences).

### Analyses and visualization of inter-hemispheric asymmetries

As the main aim of this study was to characterize the inter-hemispheric asymmetries of folding patterns, for each proxy, we focused on the dimensions discriminating left and right hemispheres, out of the N_dim_ dimensions capturing the shape variability. Only dimensions showing a statistically significant Wilcoxon signed-rank test between left and right folding pattern proxies (after Bonferroni correction) are detailed. For the analysis of the Sylvian fissure (1), the Wilcoxon signed-rank tests were applied separately to the 30w and 40w PMA proxies; we additionally tested for Spearman correlations between the positions of 30w PMA and 40w PMA proxies on the dimensions that appeared to be asymmetric at either 30w or 40w PMA.

Once the asymmetrical dimensions from the Isomap algorithm were identified, the visualization of the resulting shape features were obtained through the means of moving averages, which capture the local average shape of sulci at ten equidistant locations along the dimension (for details on their construction, see de Vareilles et al., 2021). For the longitudinal study of the Sylvian fissure (1), for visualization purposes, moving averages were generated independently for 30w and 40w PMA proxies even though the Isomap dimension of interest was computed considering the two ages together. Moreover, to highlight the difference of distribution between left and right sulcal items, we computed graphs of left versus right hemisphere distribution along the dimension, through the means of kernel density estimates. An example of sulcal projection with corresponding moving averages at both ages is presented on Figure 4.A for the Sylvian fissure, with a sample of left versus right hemisphere distribution graph at 40w PMA on Fig 4.B. In the following, only moving averages and distribution graphs (either for the 30w and 40w PMA proxies or for the left and right hemisphere proxies) are presented to describe the isomap dimensions.

**Figure 4.**
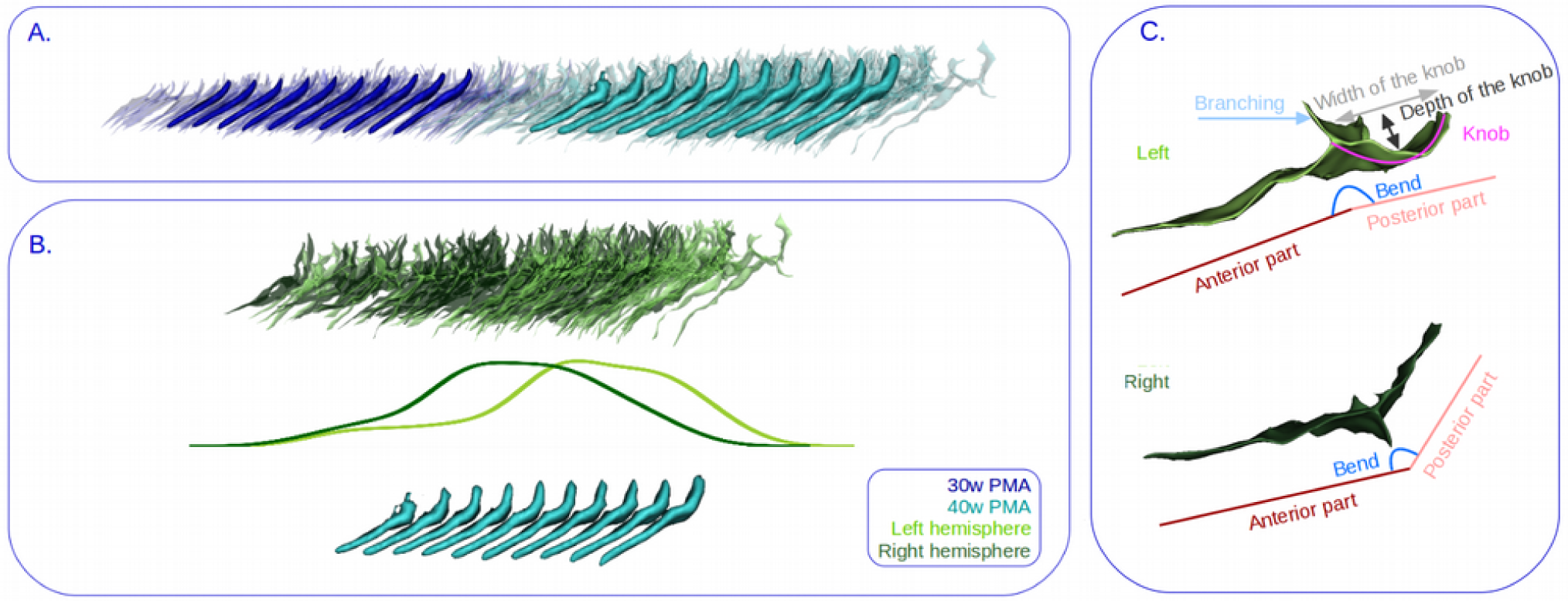
Example of folding pattern proxies and isomap representations, with reading keys for shape interpretation. A) Visualization of the first isomap dimension of the Sylvian fissure. Transparent: sulcal projections. Opaque: corresponding moving averages. B) At 40w PMA, representation of the left versus right hemisphere proxies distribution. Top: sulcal projection, Middle: left versus right hemisphere distribution graph, Bottom: moving averages. Deep blue: 30w PMA, cyan: 40w PMA, light green: left hemisphere, Dark green: right hemisphere. C) Reading keys for the shape of the Sylvian fissure, presented on Top: a 40w PMA left Sylvian fissure and on Bottom: a 40w PMA right Sylvian fissure after mirror-flip. The posterior part may (as in the example of left fissure) or may not (as in the example of flipped right fissure) contain a knob, which itself may occupy all of the posterior part or only part of it.

### Analysis of inter-region shape correlations

In order to assess whether the proxies’ patterns captured in the different sulcal regions tended to match, we computed the Spearman correlations between the Isomap positions of the 40w PMA proxies for the Sylvian fissure, the perisylvian region, and the inferior frontal gyrus. The p-values obtained were corrected according to the Bonferroni method.

## Results

### Longitudinal analysis of the Sylvian fissure shape characteristics

The Sylvian fissure’s shape variability was investigated by considering the 30w and 40w PMA proxies together. Out of the 6 dimensions of interest, 3 showed asymmetries at both ages, and the 3 others showed asymmetries only at one age (cf. Sup. Table S2.A). Actually, the 3 dimensions showing consistent asymmetries across development showed significant correlations between the positioning of the 30w and 40w PMA proxies on both hemispheres (cf. Sup. Table S3). A reading key for the shape description of the Sylvian fissure is presented on Figure 4.C, and the 6 isomap dimensions obtained on the Sylvian fissure are represented on Figure 5. At both ages, left Sylvian fissures tend to be longer, flatter or even bending backwards, and to branch more than right Sylvian fissures. Additionally, compared to the right fissures, the left Sylvian fissures at 30w PMA showed a narrower knob in the posterior part and a shorter anterior part relatively to the posterior part. At 40w PMA exclusively, the left Sylvian fissures showed a deeper knob than their right hemisphere counterparts.

**Figure 5.**
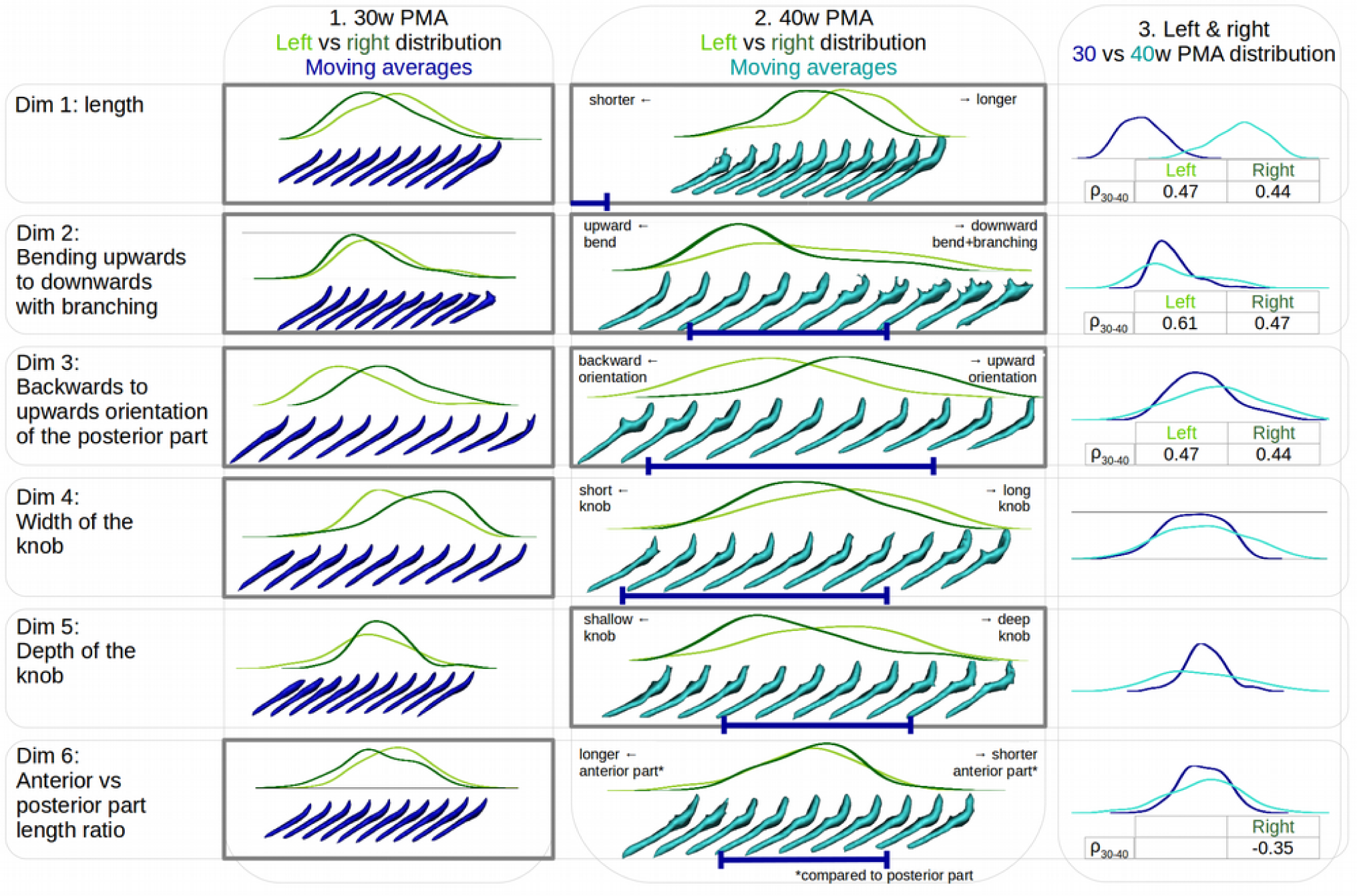
Description of each of the 6 isomap dimensions obtained on the Sylvian fissure. Left versus right hemisphere distribution plots of the Sylvian fissure are represented at 30w PMA (1) and 40w PMA (2), along with the respective moving averages. On the 40w PMA column (2), a deep blue interval is indicated under the 40w PMA moving averages to locate the span of the corresponding 30w PMA moving averages. 30w versus 40w PMA distribution plots are further superposed, combining left and right proxies of the Sylvian fissures (3). In each column, grey bold contours indicate dimensions showing a statistically significant left vs right hemisphere proxies’ positioning (for 1 and 2, see Sup. Table S2). When significant, the hemisphere-specific Spearman correlation (ρ_30-40_) is indicated under the 30w versus 40w PMA position comparisons (3, see Sup. Table S3). Deep blue: relative to 30w PMA proxies, cyan: relative to 40w PMA proxies, dark green: relative to right hemisphere proxies, light green: relative to left hemisphere proxies.

### Shape asymmetries of the perisylvian region at term-equivalent age

Out of the 10 dimensions relevant to describe the shape characteristics of the perisylvian region, only the first one was asymmetric, but very significantly (cf Sup. Table S2.B). Together with a reading key (Fig 6.A), this dimension is represented on Figure 6.B illustrating that 1) left proxies show a higher density of sulci in the inferior frontal region than right ones, 2) left proxies have a narrower gap between the frontal and temporal lobes than right proxies, and 3) the temporo-parietal region of left proxies is less curvy than that of right ones. Incidentally, right perisylvian region proxies seem to have been drawn from hemispheres with curvier Sylvian fissures while left ones from hemispheres with flatter Sylvian fissures. This suspicion was visually confirmed by projecting the individual Sylvian fissures on this perisylvian region’s isomap dimension and generating the resulting moving averages (Fig 6.C): the dimension seems to capture a shift from an upward to a backward orientation of the Sylvian fissure.

**Figure 6.**
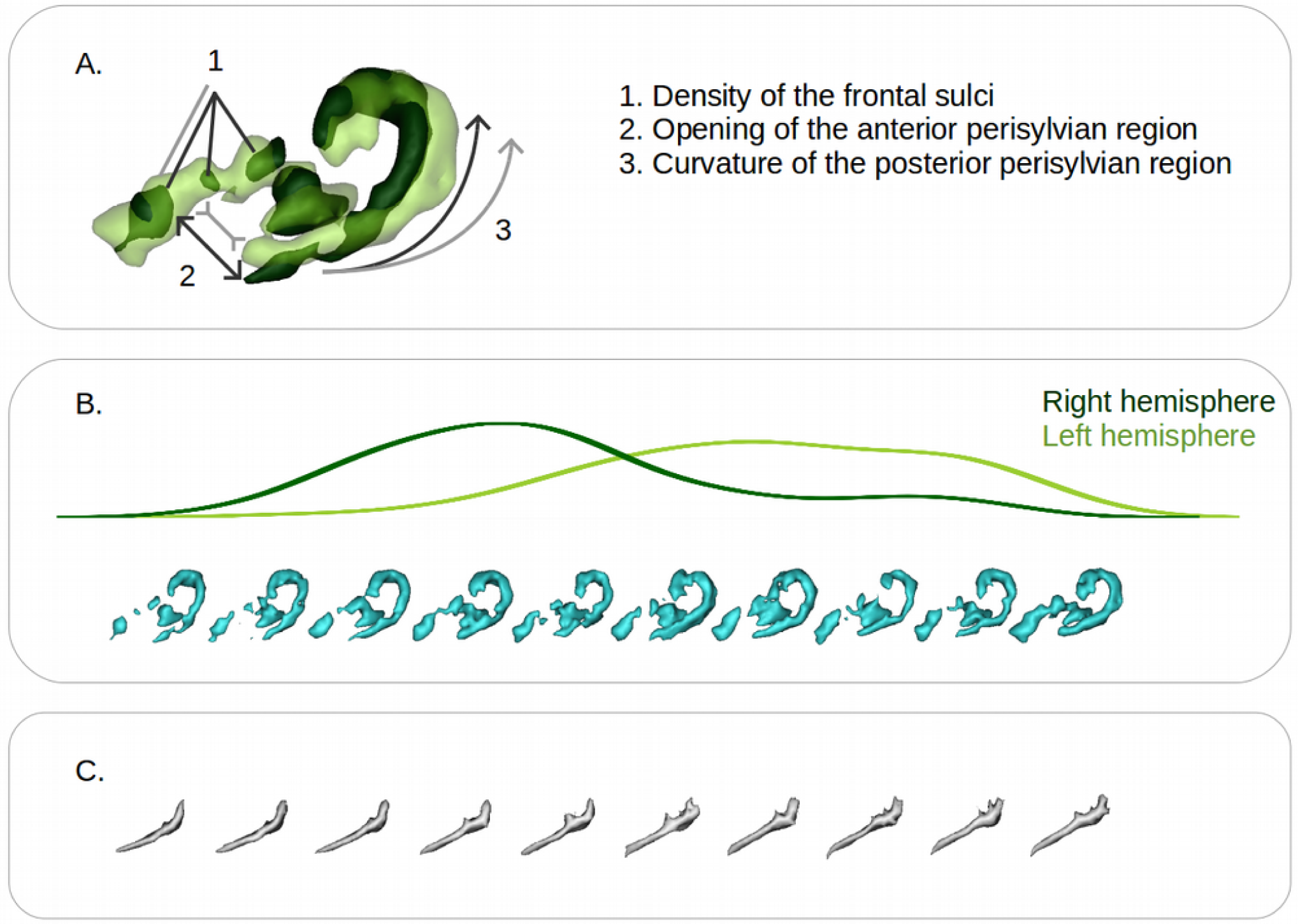
Representation of the first isomap dimension obtained for the perisylvian region at 40w PMA. A. Reading key for the perisylvian region, based on the projection of the two extreme moving averages (representative of the right hemisphere: dark green; representative of left hemisphere: light green) and illustration of their main differences, illustrative of the shape features captured (dark grey, representative of the right hemisphere, light grey, representative of the left hemisphere). B. Left (light green) versus right (dark green) hemisphere distribution plot of the perisylvian folding pattern proxies along with the corresponding moving averages (cyan). C. Moving averages of the Sylvian fissures obtained by projection on the perisylvian region isomap dimension.

This was also confirmed by the significant negative correlation (cf Sup. Table S4) between the perisylvian region’s dimension and the 40w PMA Sylvian fissure’s 1^st^ (length), 3^rd^ (backwards to upwards orientation of the posterior part), and 5^th^ (depth of the supra-marginal knob) dimensions. The highest correlation was captured on the 3^rd^ dimension.

### Shape asymmetries of the inferior frontal gyrus at term-equivalent age

Again, out of the 8 dimensions highlighted in the analysis of the inferior frontal gyrus shape characteristics, only the first dimension showed a statistically highly significant hemispheric asymmetry (cf Sup. Table S2.C), represented on Figure 7.

**Figure 7.**
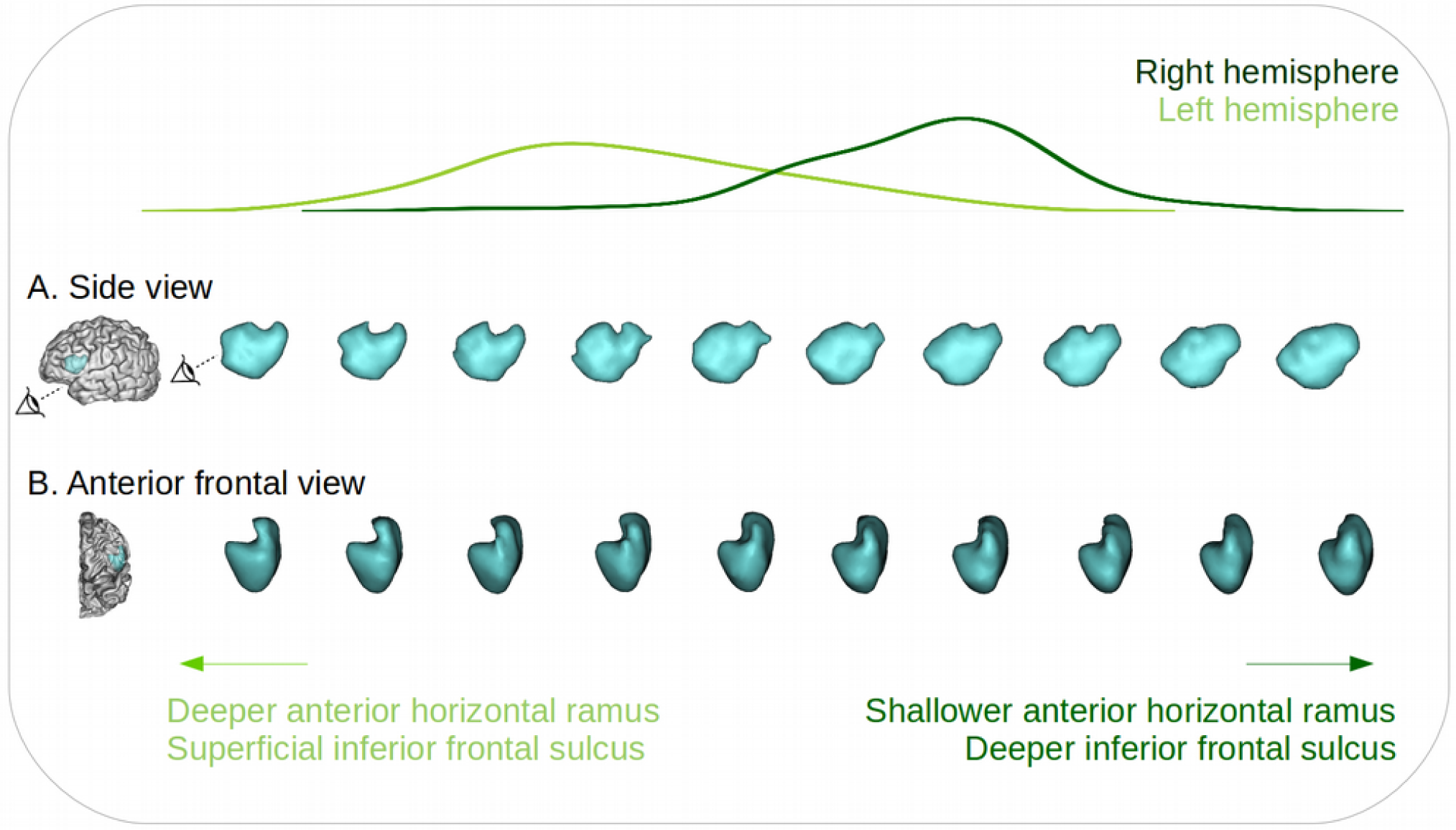
Representation of the first isomap dimension obtained for the inferior frontal gyrus at 40w PMA. Left (light green) versus right (dark green) hemisphere distribution plot of the distribution of the inferior frontal gyrus proxies, along with the corresponding moving averages, from a sideways view (A) and an anterior front-side view (B). Two eye indicators are represented in (A) to illustrate the orientation of the anterior frontal view (B).

The shape characteristics of this isomap dimension were quite complex to interpret probably because of the high inter-individual variability. Nevertheless, on the side view of this dimension, we observed a hole in the superior part of the moving averages which was predominant in the left hemisphere compared to the right. More importantly, on the anterior frontal view, we observed that the left hemisphere proxies presented a deeper inferior part of the moving average – capturing the anterior horizontal ramus – and a thinner upper part – capturing the anterior part of the inferior frontal sulcus – than their right counterpart, along with a different angle between the inferior and upper part of the moving averages, switching from acute on the left hemisphere to perpendicular or even slightly obtuse on the right one.

Comparing the inferior frontal region’s and the Sylvian fissure’s Isomap dimensions, we found statistically significant correlations for the first 3 dimensions of the fissure (cf Sup. Table S4), the strongest one being with the 3^rd^ dimension once again, encoding for the backward to upward orientation of the posterior part of the fissure. A significant negative correlation was also observed between the inferior frontal and perisylvian regions’ Isomap dimensions (ρ = -0.47, p-value = 5.10^−9^).

## Discussion

In this study, we investigated the early shape asymmetries of the Sylvian fissure, the perisylvian region, and the inferior frontal gyrus, in a cohort of extremely preterm infants. The Sylvian fissure was investigated in a longitudinal manner, enabling us to qualify and quantify its early shape at 30w PMA and to evaluate the shape evolution dynamics until the age of normal-term birth. We then captured asymmetries in multiple interpretable shape features, and we could assess how early and how linearly they develop between 30w and 40w PMA. Around TEA, the analysis of the perisylvian region unveiled an important shape asymmetry between the hemispheres, which we suggest is due to a more pronounced opercularization on the left hemisphere compared to the right at this age. Focusing on the inferior frontal gyrus, we captured a novel strong hemispheric asymmetry that would deserve interest to better understand the development of functional lateralization for language processing.

### Methodological considerations

Before detailing the different findings of this study and their implications, we first discuss some methodological choices. The scientific opportunity of studying the early development of the 30w PMA Sylvian fissure led us to necessary arrangements for the definition of its proxy. We had to exclude the anterior part of the Sylvian fissure (anterior to the junction between the Sylvian fissure and its anterior branches) and the circular sulcus of the insula because the limited progress of opercularization at 30w PMA prevented us from a reliable segmentation for longitudinal comparison.

We kept this focus in mind when choosing the sulci to preselect for the perisylvian region proxy at around TEA. We selected all the sulci neighboring the Sylvian fissure from the frontal, parietal and temporal lobes, except for the sulci from the extreme anterior frontal and temporal cortices, then excluding the orbitofrontal sulci and the polar temporal sulcus, in order to focus on the lateral view of the perisylvian region, and to focus on objects neighboring the Sylvian fissure proxy, which did not extend anteriorly. Because of the variability of sulcal configurations represented in this cohort, we preferred to preselect a list of sulci and weight their inclusion by their distance to the Sylvian fissure rather than including them whole. This allowed us to ensure that even in chaotic circumstances, we would efficiently capture the relevant sulci (close to the Sylvian fissure), and that irrelevant sulcal elements would be sufficiently weighted out to be negligible. The same method was applied to select a relevant proxy for the inferior frontal region.

In addition to the list of sulci preselected before weighting their inclusion in the study, two choices impacted our final perisylvian and inferior frontal proxies selection: the Sylvian fissure-derived items selected to weight the inclusion of sulci (the Sylvian fissure’s main branch for the perisylvian region, and the most anterior point of the Sylvian fissure for the inferior frontal region), and the choice of σ. For the perisylvian region, we chose to retain only the main part of the Sylvian fissure -and not the branch posterior to the central sulcus for example-in order to select an anatomical object with the lowest inter-individual variability. In the inferior frontal gyrus, using only the most anterior point of the Sylvian fissure proxy was optimal to capture gyri specifically around this marker, allowing for the inferior parts of the precentral gyrus and the relevant part of the inferior frontal sulcus to both be included, and with a similar weight.

In terms of σ², its manual selection was decided in order to ensure that each selected region would capture relevant landmarks: neither too few nor too many sulci selected, which would either result in an incomplete or a too broad depiction of the two regions assessed. For example, in the perisylvian region, we aimed at selecting only the parts of the superior and inferior temporal sulci which were adjacent to the Sylvian fissure. These kinds of consideration led to a value of σ² which could be fine-tuned within a range corresponding to this type of criterion. Yet, we argue that small variations of σ² would have led to very similar proxies and therefore equivalent results, and that big variations of σ² would change the specific focus of the study.

### Asymmetries linked to the Sylvian fissure’s shape

Taking into consideration these methodological decisions, let us focus now on the findings we have reported. First of all, multiple shape features with hemispheric asymmetries were observed in the Sylvian fissure, at 30w and 40w PMA. Three shape features captured asymmetries at both ages: the left Sylvian fissure tends to be longer, flatter or curved backwards, and to branch more than its right counterpart, as soon as 30w PMA and ongoing at 40w PMA. Additionally, at 30w PMA only, the ratio between the lengths of the anterior part and the posterior part of the fissure was smaller on the left than on the right side. And at 40w PMA only, the left Sylvian fissure presented a deeper supra-marginal knob than the right one. The length asymmetry of the Sylvian fissure (along with an asymmetry in surface area which we did not quantify here) has already been reported in multiple studies, either directly through the Sylvian fissure (in the adult: Lyttleton et al., 2009; comparatively in the child, adolescent and adult: Sowell et al., 2002; in the preterm newborn: Kersbergen et al., 2016), or, for its posterior extent, through the length asymmetry of the *planum temporale*, an adjacent gyral structure (in the adult: Lyttleton et al., 2009; in the full-term newborn: Glasel et al., 2011; in the preterm newborn: Dubois et al., 2010). This length asymmetry was reported to be due to the left Sylvian fissure extending both more anteriorly and more posteriorly than its right counterpart (Dubois et al., 2010), and to increase with age (Sowell et al., 2002). The curvature asymmetry has also previously been reported in the adult (Lyttleton et al., 2009; the posterior ascending ramus being more anterior in the right hemisphere), comparatively in the child, adolescent and adult (Sowell et al., 2002), in the full-term newborn (Glasel et al., 2011), and in the fetus (Habas et al., 2012; the posterior temporal operculum being more concave in the right hemisphere). To the best of our knowledge, the tendency of the left Sylvian fissure to bend backwards and to branch more than the right one is previously unreported, all the more so in the developing brain. Nevertheless, the backward bend might be paralleled with a pattern-oriented study of the parietal opercular region (Steinmetz et al., 1990), which reported a configuration of the posterior part of the Sylvian fissure – only observed in the left hemisphere – without a posterior ascending ramus, suggesting a flat or bending backwards Sylvian fissure. Interestingly, it should be noted that these three shape features capturing asymmetries at both 30 and 40w PMA are the only ones that showed significant positive correlations between the two ages, suggesting an early asymmetry encoding and a development according to the early shape predisposition.

We further observed two asymmetries encoded only at 30w PMA: the width of the knob, and the ratio between the anterior part of the Sylvian fissure and its posterior part. These seem to encode transient shape encoding differences and we suggest that they might reflect differences in developmental trajectories between left and right hemisphere Sylvian fissures. At 40w PMA only, an asymmetry was captured in the depth of the knob of the posterior part of the Sylvian fissure. This knob is located under the supramarginal gyrus, and its deepening in the left hemisphere is most likely due to a bigger supramarginal gyrus in the left hemisphere, which has been reported repeatedly in the adult (Lyttleton et al., 2009; Kong et al., 2018) and infant brain (Li et al., 2014). Interestingly, this observation is consistent with the fact that, in the nomenclature we use, the only sulcus which is not defined bilaterally but only in the left hemisphere is specific to the supramarginal gyrus (Perrot et al., 2011; Borne et al., 2020). Not capturing this asymmetry at 30w PMA suggests that the differential development of the supramarginal gyrus between the two hemispheres is progressing between 30w and 40w PMA.

Visually, the shape asymmetries of the Sylvian fissure in terms of curvature and length in its posterior region seem to have overlapped on those captured in the perisylvian region: right Sylvian fissures are more curvy and shorter than their left counterparts, and these same curvature and posterior extension seem to be captured in the perisylvian region. This was confirmed by the significant correlations captured between the perisylvian and three of the Sylvian fissure Isomap dimensions. This might rely on the fact that the opercularization process, leading to the Sylvian fissure formation, also displaces all the surrounding folding regions. This mix between a less curved and more posteriorly extended left perisylvian region is compatible with a previous study which reported increased distances in the left hemisphere between the bottom point of the central sulcus and both of the posterior extremities of the Sylvian fissure and the superior temporal sulcus in the full-term infant (Glasel et al, 2011).

While we could visually expect the perisylvian dimension to correlate with the Sylvian fissure’s shape, it was not specifically the case for the inferior frontal region, whose pattern should *a priori* not be specifically impacted by the shape of the Sylvian fissure. Yet, we observed significant correlations with 3 of its dimensions, which happened to be dimensions showing asymmetrical configurations at 40w PMA, as well as with the perisylvian region isomap dimension. The highest correlation was found with a dimension encoding for the orientation of the posterior part of the Sylvian fissure, which, aside from its link to the opercularization process, would not seem to be related to the sulcal configuration in the inferior frontal region. These correlations suggested a consistent encoding of asymmetrical shape features in the three regions.

### Opercularization is more pronounced in the left hemisphere at TEA

In the perisylvian region, apart from the previous asymmetric shape features derived from the Sylvian fissure, two additional characteristics were captured: the size of the gap between the frontal and anterior temporal sulci (smaller on the left), and the density of sulci in the frontal region (higher on the left). The volume of the part corresponding to the inferior frontal sulci appeared to be greater on the configurations corresponding to left rather than right hemispheres, even though the trend was not linear along the axis. We double-checked if this could be due to more developed inferior frontal sulci in the left hemisphere, but we found no surface asymmetry between hemispheres within the combined Sylvian fissure’s anterior branches and inferior frontal sulcus. Therefore, these characteristics pointed towards a more pronounced opercularization in the left hemisphere compared to the right one, since it means bringing closer together the frontal and temporal lobes and consequently bringing closer to the Sylvian fissure both the superior temporal sulcus and the sulci of the inferior frontal lobe. With the retained methodology, this would have induced a greater inclusion of inferior frontal and superior temporal sulci, matching the trends captured in the perisylvian region moving averages. Incidentally, we interpreted the greater consistency in capturing more sulci in the frontal lobe, relative to temporal lobe, as being due to the organization of inferior frontal sulci, globally perpendicular to the Sylvian fissure, while the superior temporal sulcus is globally parallel to the Sylvian fissure. Therefore, using our methodology, even a small advance of the inferior frontal lobe towards the insula would have induced more sulcal elements to be captured, while a small advance of the temporal lobe would have led to weight slightly more the inclusion of the superior temporal sulcus but not to capture additional sulcal elements. Therefore, the observation of asymmetries in the perisylvian proxy support the hypothesis of a more advanced or more pronounced opercularization process in the left hemisphere at TEA.

The asymmetry captured in the inferior frontal region proxy is relatively coherent with these findings as it seems to capture a more prominent anterior part in the left compared with the right hemisphere, corresponding to the horizontal ramus of the Sylvian fissure being buried deeper in the frontal cortex in the anterior frontal view. Surprisingly, contrarily to the perisylvian proxy, the inferior frontal region moving averages did not specifically capture more inferior frontal sulci on the left hemisphere. Yet, it should be reminded that both the point cloud selection and the σ parameter were different in the inferior frontal proxy: sulcal elements were retained based on their distance to a single point in the Sylvian fissure rather than their distance to the main branch, and σ was higher inducing a more global moving average shape, which may have pruned out this variability. Therefore, the methodology retained for the inferior frontal region may have dimmed out some of the sulcal variability linked to opercularization. Nevertheless, the shape asymmetry captured in the inferior frontal gyrus proxy, suggesting more deeply buried anterior rami of the Sylvian fissure, was supported by the observation that the anterior region of the Sylvian fissure (bordering the inferior frontal gyrus) seemed larger on the left hemisphere using voxel-based analyses on preterm infants between 27 and 36w PMA (Dubois et al., 2010). Our method proved the presence of a leftward asymmetry in this region which was previously inconsistently reported (Sprung-Much et al., 2021). This asymmetry could be fundamental for the development of language lateralization since this is the anatomical region corresponding to Broca’s area.

### Opercularization and sulcation show different dynamics

This more pronounced opercularization on the left hemisphere at TEA contrasts with the previously reported rightward advance in sulcation during pregnancy and around birth (Chi et al., 1977; Dubois et al., 2008; Habas et al., 2012) and led us to question the difference in dynamics between the two folding processes. The strong inter-hemispheric asymmetries captured in the Sylvian fissure, highly reproducible within the human population and reported as soon as 23w GA (Habas et al., 2012), suggest an important genetic control over opercularization, which is supported by the important asymmetry captured in the perisylvian and inferior frontal regions in the present study. Incidentally, asymmetrically expressed genes have been identified specifically in perisylvian cortices (Sun et al., 2005) and linked to the asymmetry of the planum temporale (Carion-Castillo et al., 2020). Moreover, the gene expression profile of the pre-folded brain has been reported to be significantly different in the developing opercular cortex compared to the insular cortex (Mallela et al., 2020).

Interestingly, another folding asymmetry is frequently observed in the developing brain, in the superior temporal sulcus’s depth (Dubois et al., 2008; Glasel et al., 2011; Habas et al., 2012; Leroy et al., 2015), yet it has been reported to be uncorrelated to the Sylvian fissure’s asymmetry (Glasel et al., 2011). This could be due to a different genetic control over these two areas, which would corroborate a difference in dynamics between opercularization and sulcation.

Incidentally, after observing the difference in dynamics between the two processes, one can wonder whether the whole Sylvian fissure is due to opercularization or if the most posterior part, posterior to the *planum temporale* and extending within the parietal cortex, as defined in Altarelli et al., 2014, may be due to sulcation. The fact that the 4^th^ dimension, encoding the length of the supra-marginal knob, showed a higher (and almost significant) correlation between 30w and 40w PMA in the right hemisphere than in the left, may point towards the latter hypothesis. Indeed, this could be related to the prenatal sulcation advance in the right hemisphere, leading to an already encoded length of the intra-parietal part of the Sylvian fissure on the right but not on the left hemisphere at 30w PMA, leading to higher consistency in the encoding of this shape feature between 30 and 40w PMA on the right hemisphere.

### Perspectives

This opercularization advance was foreshadowed by previous studies. The study reporting a more extended anterior and posterior Sylvian fissure in the left hemisphere in the preterm suggested that it might either capture a developmental asynchrony or an intrinsically different structural morphology between left and right Sylvian fissure development (Dubois et al., 2010). Both seemed to be the case in the present study. In a study comparing maturation indices hemisphere-wise along a set of sulci around the Sylvian fissure, the most leftward asymmetries were captured in the anterior precentral cortex and in the *pars opercularis* and may have captured a maturational advance linked to opercularization, even though they did not reach statistical significance (Leroy et al., 2011). Studying the maturation of opercular cortices in larger cohorts might therefore allow us to capture maturational asymmetries. Microstructural investigations have shown more consistent leftward asymmetry tendency in the inferior frontal region corresponding to Broca’s area than macroscopical investigations (Sprung-Much et al., 2021). Moreover, based on a human fetal brain atlas between 21 and 38w GA, a recent study demonstrated the link between differential genetic expression in the opercular cortices compared to the insular cortex, suggesting an important genetic control over opercularization (Mallela et al., 2020). The asynchronous development of opercularization could therefore be linked to an asymmetrical genetic regulation between hemispheres. Therefore, microstructural and genetic investigations should be undertaken in the context of opercularization in order to quantify its specificities, since its dynamics seem different from those of sulcation.

Alternatively, the current study focused on preterm infants to investigate early folding development, yet the observations at 40w PMA could and should be reproduced on term-born neonates to assess normal folding development, since extreme prematurity is associated to pathology and atypical cortical development, including in sulcation (Shimony et al., 2016; Dubois et al., 2019). In the preterm, the relationship between opercularization dynamics, prematurity, and anatomo-functional links to language should also be investigated further, as prematurity has been reported to affect both the lateralization of language (in neonates: Kwon et al., 2015; in adolescents: Scheinost et al., 2015; in adults: Tseng et al., 2019) and language scores (in children: Barnes-Davis et al., 2020; in adolescents: Scheinost et al., 2015; in adults: Tseng et al., 2019). In these studies, the only reported link to anatomy was not related to altered brain folding, but to a significant decrease in callosal connectivity accompanied by an increase in non-callosal connectivity; this non-callosal connectivity was positively correlated to language assessments in children born preterm but not in controls (Barnes-Davis et al., 2020). Language scores were additionally reported to correlate with language lateralization specifically in the population which was born very preterm (Scheinost et al., 2015), suggesting developmental adversities affecting jointly lateralization and efficiency of language networks. To assess links between folding development and language scores, an analysis between numerous sulcal metrics in the Sylvian fissure and superior temporal sulcus (area, depth, and sulcal index) in the present cohort reported significant correlations to receptive language outcome assessed at around 2 years of age (Kersbergen et al., 2016).

Therefore, to complement our study, it would be most informative to investigate further the specificities of opercularization, either through the observation of pattern dynamics at more time-points, through maturational or microstructural considerations, or through the evaluation of asymmetrical genetic regulation. Our method should further be applied to control term-born neonates to quantify the extent to which observations of early asymmetries in perisylvian regions in preterm infants correspond to a typical variability. A functional investigation would also be informative to assess whether the fine shape of any of the three proxies that we have investigated relates to later language outcomes.

## Acknowledgements

The authors thank Y. Leprince, L. Perus, L. Guillon and W. Shu-Quartier-Dit-Maire for their participation in discussions about this study.

## References

Altarelli, I., Leroy, F., Monzalvo, K., Fluss, J., Billard, C., Dehaene-Lambertz, G., Galaburda, A.M., Ramus, F., 2014. Planum temporale asymmetry in developmental dyslexia: Revisiting an old question: Planum Temporale Asymmetry in Dyslexia. Hum. Brain Mapp. 35, 5717–5735. https://doi.org/10.1002/hbm.22579

Amiez, C., Sallet, J., Novek, J., Hadj-Bouziane, F., Giacometti, C., Andersson, J., Hopkins, W.D., Petrides, M., 2021. Chimpanzee histology and functional brain imaging show that the paracingulate sulcus is not human-specific. Commun Biol 4, 54. https://doi.org/10.1038/s42003-020-01571-3

Barnes-Davis, M.E., Williamson, B.J., Merhar, S.L., Holland, S.K., Kadis, D.S., 2020. Rewiring the extremely preterm brain: Altered structural connectivity relates to language function. NeuroImage: Clinical 25, 102194. https://doi.org/10.1016/j.nicl.2020.102194

Besl, P.J., McKay, N., 1992. A method for registration of 3-D shapes. IEEE Transactions on Pattern Analysis and Machine Intelligence 14, 239–256. https://doi.org/10.1109/34.121791

Borne, L., 2019. Design of a top-down computer vision algorithm dedicated to the recognition of cortical sulci. PhD thesis, Université Paris-Saclay.

Borne, L., Rivière, D., Mancip, M., Mangin, J.-F., 2020. Automatic labeling of cortical sulci using patch-or CNN-based segmentation techniques combined with bottom-up geometric constraints. Medical Image Analysis 62, 101651. https://doi.org/10.1016/j.media.2020.101651

Bouyssi-Kobar, M., Murnick, J., Tinkleman, L., Robertson, R.L., Limperopoulos, C., 2016. Third Trimester Brain Growth in Preterm Infants Compared With In Utero Healthy Fetuses 138, 13.

Carrion-Castillo, A., Pepe, A., Kong, X.-Z., Fisher, S.E., Mazoyer, B., Tzourio-Mazoyer, N., Crivello, F., Francks, C., 2020. Genetic effects on planum temporale asymmetry and their limited relevance to neurodevelopmental disorders, intelligence or educational attainment. Cortex 124, 137–153. https://doi.org/10.1016/j.cortex.2019.11.006

Chi, J.G., Dooling, E.C., Gilles, F.H., 1977. Gyral development of the human brain. Ann Neurol. 1, 86–93. https://doi.org/10.1002/ana.410010109

de Vareilles, H., Rivière, D., Sun, Z., Fischer, C., Leroy, F., Neumane, S., Stopar, N., Eijsermans, R., Ballu, M., Tataranno, M., Benders, M., Mangin, J., Dubois, J., 2022. Shape variability of the central sulcus in the developing brain: a longitudinal descriptive and predictive study in preterm infants. NeuroImage 118837. https://doi.org/10.1016/j.neuroimage.2021.118837

Dubois, J., Benders, M., Cachia, A., Lazeyras, F., Ha-Vinh Leuchter, R., Sizonenko, S.V., Borradori-Tolsa, C., Mangin, J.F., Huppi, P.S., 2008. Mapping the Early Cortical Folding Process in the Preterm Newborn Brain. Cerebral Cortex 18, 1444–1454. https://doi.org/10.1093/cercor/bhm180

Dubois, J., Benders, M., Lazeyras, F., Borradori-Tolsa, C., Leuchter, R.H.-V., Mangin, J.F., Hüppi, P.S., 2010. Structural asymmetries of perisylvian regions in the preterm newborn. NeuroImage 52, 32–42. https://doi.org/10.1016/j.neuroimage.2010.03.054

Dubois, J., Lefèvre, J., Angleys, H., Leroy, F., Fischer, C., Lebenberg, J., Dehaene-Lambertz, G., Borradori-Tolsa, C., Lazeyras, F., Hertz-Pannier, L., Mangin, J.-F., Hüppi, P.S., Germanaud, D., 2019. The dynamics of cortical folding waves and prematurity-related deviations revealed by spatial and spectral analysis of gyrification. NeuroImage 185, 934–946. https://doi.org/10.1016/j.neuroimage.2018.03.005

Glasel, H., Leroy, F., Dubois, J., Hertz-Pannier, L., Mangin, J.F., Dehaene-Lambertz, G., 2011. A robust cerebral asymmetry in the infant brain: The rightward superior temporal sulcus. NeuroImage 58, 716–723. https://doi.org/10.1016/j.neuroimage.2011.06.016

Habas, P.A., Scott, J.A., Roosta, A., Rajagopalan, V., Kim, K., Rousseau, F., Barkovich, A.J., Glenn, O.A., Studholme, C., 2012. Early Folding Patterns and Asymmetries of the Normal Human Brain Detected from in Utero MRI. Cerebral Cortex 22, 13–25. https://doi.org/10.1093/cercor/bhr053

Hickok, G., Poeppel, D., 2007. The cortical organization of speech processing. Nat Rev Neurosci 8, 393–402. https://doi.org/10.1038/nrn2113

Hill, J., Dierker, D., Neil, J., Inder, T., Knutsen, A., Harwell, J., Coalson, T., Van Essen, D., 2010. A Surface-Based Analysis of Hemispheric Asymmetries and Folding of Cerebral Cortex in Term-Born Human Infants. Journal of Neuroscience 30, 2268–2276. https://doi.org/10.1523/JNEUROSCI.4682-09.2010

Hou, L., Xiang, L., Crow, T.J., Leroy, F., Rivière, D., Mangin, J.-F., Roberts, N., 2019. Measurement of Sylvian fissure asymmetry and occipital bending in humans and Pan troglodytes. NeuroImage 184, 855–870. https://doi.org/10.1016/j.neuroimage.2018.08.045

Jiang, X., Zhang, T., Zhang, S., Kendrick, K.M., Liu, T., 2021. Fundamental functional differences between gyri and sulci: implications for brain function, cognition, and behavior. Psychoradiology 1, 23–41. https://doi.org/10.1093/psyrad/kkab002

Kersbergen, K.J., Leroy, F., Išgum, I., Groenendaal, F., de Vries, L.S., Claessens, N.H.P., van Haastert, I.C., Moeskops, P., Fischer, C., Mangin, J.-F., Viergever, M.A., Dubois, J., Benders, M.J.N.L., 2016. Relation between clinical risk factors, early cortical changes, and neurodevelopmental outcome in preterm infants. NeuroImage 142, 301–310. https://doi.org/10.1016/j.neuroimage.2016.07.010

Knecht, S., Dräger, B., Deppe, M., Bobe, L., Lohmann, H., Flöel, A., Ringelstein, E.-B., Henningsen, H., 2000. Handedness and hemispheric language dominance in healthy humans. Brain 123, 2512–2518. https://doi.org/10.1093/brain/123.12.2512

Kong, X.-Z., Mathias, S.R., Guadalupe, T., ENIGMA Laterality Working Group, Glahn, D.C., Franke, B., Crivello, F., Tzourio-Mazoyer, N., Fisher, S.E., Thompson, P.M., Francks, C., 2018. Mapping cortical brain asymmetry in 17,141 healthy individuals worldwide via the ENIGMA Consortium. Proc Natl Acad Sci USA 115, E5154–E5163. https://doi.org/10.1073/pnas.1718418115

Kostović, I., Sedmak, G., Judaš, M., 2019. Neural histology and neurogenesis of the human fetal and infant brain. NeuroImage 188, 743–773. https://doi.org/10.1016/j.neuroimage.2018.12.043

Kwon, S.H., Scheinost, D., Lacadie, C., Sze, G., Schneider, K.C., Dai, F., Constable, R.T., Ment, L.R., 2015. Adaptive mechanisms of developing brain: Cerebral lateralization in the prematurely-born. NeuroImage 108, 144–150. https://doi.org/10.1016/j.neuroimage.2014.12.032

Le Guen, Y., Leroy, F., Philippe, C., IMAGEN consortium, Mangin, J.-F., Dehaene-Lambertz, G., Frouin, V., 2019. Enhancer locus in ch14q23.1 modulates brain asymmetric temporal regions involved in language processing (preprint). Neuroscience. https://doi.org/10.1101/539189

Lefèvre, J., Germanaud, D., Dubois, J., Rousseau, F., de Macedo Santos, I., Angleys, H., Mangin, J.-F., Hüppi, P.S., Girard, N., De Guio, F., 2016. Are Developmental Trajectories of Cortical Folding Comparable Between Cross-sectional Datasets of Fetuses and Preterm Newborns? Cereb. Cortex 26, 3023–3035. https://doi.org/10.1093/cercor/bhv123

Leroy, F., Glasel, H., Dubois, J., Hertz-Pannier, L., Thirion, B., Mangin, J.-F., Dehaene-Lambertz, G., 2011. Early Maturation of the Linguistic Dorsal Pathway in Human Infants. Journal of Neuroscience 31, 1500–1506. https://doi.org/10.1523/JNEUROSCI.4141-10.2011

Leroy, F., Cai, Q., Bogart, S.L., Dubois, J., Coulon, O., Monzalvo, K., Fischer, C., Glasel, H., Van der Haegen, L., Bénézit, A., Lin, C.-P., Kennedy, D.N., Ihara, A.S., Hertz-Pannier, L., Moutard, M.-L., Poupon, C., Brysbaert, M., Roberts, N., Hopkins, W.D., Mangin, J.-F., Dehaene-Lambertz, G., 2015. New human-specific brain landmark: The depth asymmetry of superior temporal sulcus. Proc Natl Acad Sci USA 112, 1208–1213. https://doi.org/10.1073/pnas.1412389112

Li, G., Wang, L., Shi, F., Lyall, A.E., Lin, W., Gilmore, J.H., Shen, D., 2014. Mapping Longitudinal Development of Local Cortical Gyrification in Infants from Birth to 2 Years of Age. Journal of Neuroscience 34, 4228–4238. https://doi.org/10.1523/JNEUROSCI.3976-13.2014

Llinares-Benadero, C., Borrell, V., 2019. Deconstructing cortical folding: genetic, cellular and mechanical determinants. Nat Rev Neurosci 20, 161–176. https://doi.org/10.1038/s41583-018-0112-2

Lyttelton, O.C., Karama, S., Ad-Dab’bagh, Y., Zatorre, R.J., Carbonell, F., Worsley, K., Evans, A.C., 2009. Positional and surface area asymmetry of the human cerebral cortex. NeuroImage 46, 895–903. https://doi.org/10.1016/j.neuroimage.2009.03.063

Mallela, A.N., Deng, H., Brisbin, A.K., Bush, A., Goldschmidt, E., 2020. Sylvian fissure development is linked to differential genetic expression in the pre-folded brain. Sci Rep 10, 14489. https://doi.org/10.1038/s41598-020-71535-4

Mangin, J.-F., Frouin, V., Bloch, I., Régis, J., López-Krahe, J., 1995. From 3D magnetic resonance images to structural representations of the cortex topography using topology preserving deformations. J Math Imaging Vis 5, 297–318. https://doi.org/10.1007/BF01250286

Mangin, J.-F., Auzias, G., Coulon, O., Sun, Z.Y., Rivière, D., Régis, J., 2015. Sulci as Landmarks, in: Brain Mapping. Elsevier, pp. 45–52. https://doi.org/10.1016/B978-0-12-397025-1.00198-6

Moeskops, P., Benders, M.J.N.L., Kersbergen, K.J., Groenendaal, F., de Vries, L.S., Viergever, M.A., Išgum, I., 2015. Development of Cortical Morphology Evaluated with Longitudinal MR Brain Images of Preterm Infants. PLoS ONE 10, e0131552. https://doi.org/10.1371/journal.,pone.0131552

Perrot, M., Rivière, D., Mangin, J.-F., 2011. Cortical sulci recognition and spatial normalization. Medical Image Analysis 15, 529–550. https://doi.org/10.1016/j.media.2011.02.008

Rivière, D., Leprince, Y., Labra, N., Vindas, N., Foubet, O., Cagna, B., Loh, K.K., Hopkins, W., Balzeau, A., Mancip, M., Lebenberg, J., Cointepas, Y., Coulon, O., Mangin, J.-F., 2022. Browsing Multiple Subjects When the Atlas Adaptation Cannot Be Achieved via a Warping Strategy. Front. Neuroinform. 16, 803934. https://doi.org/10.3389/fninf.2022.803934

Scheinost, D., Lacadie, C., Vohr, B.R., Schneider, K.C., Papademetris, X., Constable, R.T., Ment, L.R., 2015. Cerebral Lateralization is Protective in the Very Prematurely Born. Cerebral Cortex 25, 1858–1866. https://doi.org/10.1093/cercor/bht430

Shimony, J.S., Smyser, C.D., Wideman, G., Alexopoulos, D., Hill, J., Harwell, J., Dierker, D., Van Essen, D.C., Inder, T.E., Neil, J.J., 2016. Comparison of cortical folding measures for evaluation of developing human brain. NeuroImage 125, 780–790. https://doi.org/10.1016/j.neuroimage.2015.11.001

Sowell, E.R., Thompson, P.M., Rex, D., Kornsand, D., Tessner, K.D., Jernigan, T.L., Toga, A.W., 2002. Mapping Sulcal Pattern Asymmetry and Local Cortical Surface Gray Matter Distribution In Vivo: Maturation in Perisylvian Cortices. Cerebral Cortex 12, 17–26. https://doi.org/10.1093/cercor/12.1.17

Sprung-Much, T., Eichert, N., Nolan, E., Petrides, M., 2021. Broca’s area and the search for anatomical asymmetry: commentary and perspectives. Brain Struct Funct 227, 441–449. https://doi.org/10.1007/s00429-021-02357-x

Steinmetz, H., 1990. Sulcus topography of the parietal opercular region: An anatomic and MR study. Brain and Language 38, 515–533. https://doi.org/10.1016/0093-934X(90)90135-4

Sun, T., Collura, R.V., Ruvolo, M., Walsh, C.A., 2006. Genomic and Evolutionary Analyses of Asymmetrically Expressed Genes in Human Fetal Left and Right Cerebral Cortex. Cerebral Cortex 16, i18–i25. https://doi.org/10.1093/cercor/bhk026

Sun, Z.Y., Klöppel, S., Rivière, D., Perrot, M., Frackowiak, R., Siebner, H., Mangin, J.-F., 2012. The effect of handedness on the shape of the central sulcus. NeuroImage 60, 332–339. https://doi.org/10.1016/j.neuroimage.2011.12.050

Sun, Z.Y., Pinel, P., Rivière, D., Moreno, A., Dehaene, S., Mangin, J.-F., 2016. Linking morphological and functional variability in hand movement and silent reading. Brain Struct Funct 221, 3361–3371. https://doi.org/10.1007/s00429-015-1106-8

Tenenbaum, J.B., 2000. A Global Geometric Framework for Nonlinear Dimensionality Reduction. Science 290, 2319–2323. https://doi.org/10.1126/science.290.5500.2319

Tissier, C., Linzarini, A., Allaire-Duquette, G., Mevel, K., Poirel, N., Dollfus, S., Etard, O., Orliac, F., Peyrin, C., Charron, S., Raznahan, A., Houdé, O., Borst, G., Cachia, A., 2018. Sulcal Polymorphisms of the IFC and ACC Contribute to Inhibitory Control Variability in Children and Adults. eNeuro 5, ENEURO.0197-17.2018. https://doi.org/10.1523/ENEURO.0197-17.2018

Tseng, C.-E.J., Froudist-Walsh, S., Kroll, J., Karolis, V., Brittain, P.J., Palamin, N., Clifton, H., Counsell, S.J., Williams, S.C.R., Murray, R.M., Nosarti, C., 2019. Verbal Fluency Is Affected by Altered Brain Lateralization in Adults Who Were Born Very Preterm. eNeuro 6, ENEURO.0274-18.2018. https://doi.org/10.1523/ENEURO.0274-18.2018

