## Supplementary material for "Exploring the emergence of morphological asymmetries around the brain’s Sylvian fissure: a longitudinal study of shape variability in preterm infants"

Annex 1: List of sulci preselected for the perisylvian region, ordered by lobe (except for the Sylvian fissure):

| |
|---|
| Frontal lobe: |
| Inferior frontal sulcus |
| Inferior and intermediary parts of the precentral sulcus |
| Central sulcus* |
| Parietal lobe: |
| Inferior part of the post-central sulcus |
| Intraparietal sulcus (including ramifications) |
| Sulcus of the supra-marginal gyrus |
| Temporal lobe: |
| Superior temporal sulcus (including ramifications) |
| Inferior temporal sulcus |
| Polar temporal sulcus |
| Sylvian fissure area: (sylvian fissure branches excluded from the sylvian fissure proxy) |
| Planum temporale sulcus |
| Transverse retrocentral ramus |
| Posterior sub-central ramus |
| Anterior sub-central ramus |
| Ascending ramus |
| Diagonal sulcus |
| Horizontal ramus |

*Any item which did not fit any sulci label was labeled as unknown. The unkown labeled items were included in the preselected list. *sulci between lobes were affected arbitrarily to one of its neighboring lobes*

### Supplementary Materials

Sup. Table S1. Pearson correlations between Isomap positions and PMA at MRI for the three studies (n=71)

| | r | | | |
|---|---|---|---|---|
| | *Sylvian fissure* | | *Perisylvian region* | *Inferior frontal region* |
| | At 30w PMA | At 40w PMA | At 40w PMA | At 40w PMA |
| Dimension 1 | 0.38 | 0.12 | -0.14 | -0.01 |
| Dimension 2 | 0.04 | -0.21 | 0.07 | 0.00 |
| Dimension 3 | 0.02 | -0.13 | -0.02 | 0.03 |
| Dimension 4 | -0.14 | 0.00 | 0.06 | 0.14 |
| Dimension 5 | -0.15 | -0.03 | 0.03 | 0.07 |
| Dimension 6 | 0.16 | 0.02 | 0.02 | -0.15 |
| Dimension 7 | - | - | -0.06 | -0.10 |
| Dimension 8 | - | - | 0.00 | 0.00 |
| Dimension 9 | - | - | 0.20 | - |
| Dimension 10 | - | - | 0.06 | - |

Sup. Table S2. Wilcoxon signed-rank test statistics and Bonferroni corrected p-values applied to the left versus right hemisphere isomap positions (corrected for PMA)

| | Stat (corrected p-value) | | | |
|---|---|---|---|---|
| | *A. Sylvian fissure* | | *B. Perisylvian region* | *C. Inferior frontal region* |
| | 30w PMA | 40w PMA | 40w PMA | 40w PMA |
| Dimension 1 | **558 ($2.10^{-4}$)** | **390 ($2.10^{-6}$)** | **224 ($2.10^{-8}$)** | **75 ($4.10^{-11}$)** |
| Dimension 2 | **772 (0.02)** | **629 ($1.10^{-3}$)** | 1177 (5.63) | 913 (0.29) |
| Dimension 3 | **400 ($3.10^{-6}$)** | **258 ($3.10^{-8}$)** | 1203 (6.68) | 1111 (2.71) |
| Dimension 4 | **596 ($6.10^{-4}$)** | 834 (0.07) | 993 (1.02) | 1098 (2.42) |
| Dimension 5 | 995 (0.62) | **693 ($4.10^{-3}$)** | 964 (0.72) | 939 (0.42) |
| Dimension 6 | **725 ($9.10^{-3}$)** | 1131 (2.40) | 1202 (6.63) | 1090 (2.25) |
| Dimension 7 | - | - | 1247 (8.59) | 1140 (3.43) |
| Dimension 8 | - | - | 1113 (3.44) | 1116 (2.83) |
| Dimension 9 | - | - | 1205(6.76) | - |
| Dimension 10 | - | - | 1163 (5.10) | - |

Bold values verify corrected p-value<0.05

Sup. Table S3. Spearman correlations (statistics and Bonferroni corrected p-values) between the 30 and 40w PMA isomap positions (corrected for PMA), for the sylvian fissure analysis

| | ρ (corrected p-value) | |
|---|---|---|
| | Left hemisphere | Right hemisphere |
| Dimension 1 | **0.47 (2.10$^{-4}$)** | **0.44 (8.10$^{-4}$)** |
| Dimension 2 | **0.61 (1.10$^{-7}$)** | **0.47 (2.10$^{-4}$)** |
| Dimension 3 | **0.75 (2.10$^{-13}$)** | **0.59 (5.10$^{-7}$)** |
| Dimension 4 | 0.17 (1.01) | 0.29 (0.08) |
| Dimension 5 | 0.01 (5.65) | 0.13 (1.76) |
| Dimension 6 | -0.23 (0.34) | **-0.35 (0.02)** |

Bold values verify corrected p-value<0.05

Sup. Table S4. Spearman correlations (statistics and Bonferroni corrected p-values) of the 40w PMA Isomap positions (corrected for PMA) between the sylvian fissure, perisylvian region, and inferior frontal region proxies

| Sylvian fissure | Perisylvian region | Inferior frontal region |
|---|---|---|
| Dimension 1 | **0.37 (4.10$^{-5}$)** | **-0.27 (0.006)** |
| Dimension 2 | **0.30 (0.002)** | **-0.27 (0.007)** |
| Dimension 3 | **-0.42 (1.10$^{-6}$)** | **0.41 (3.10$^{-6}$)** |
| Dimension 4 | 0.04 (3.92) | -0.18 (0.17) |
| Dimension 5 | 0.20 (0.12) | -0.16 (0.38) |
| Dimension 6 | -0.15 (0.49) | 0.13 (0.64) |

Bold values verify corrected p-value<0.05

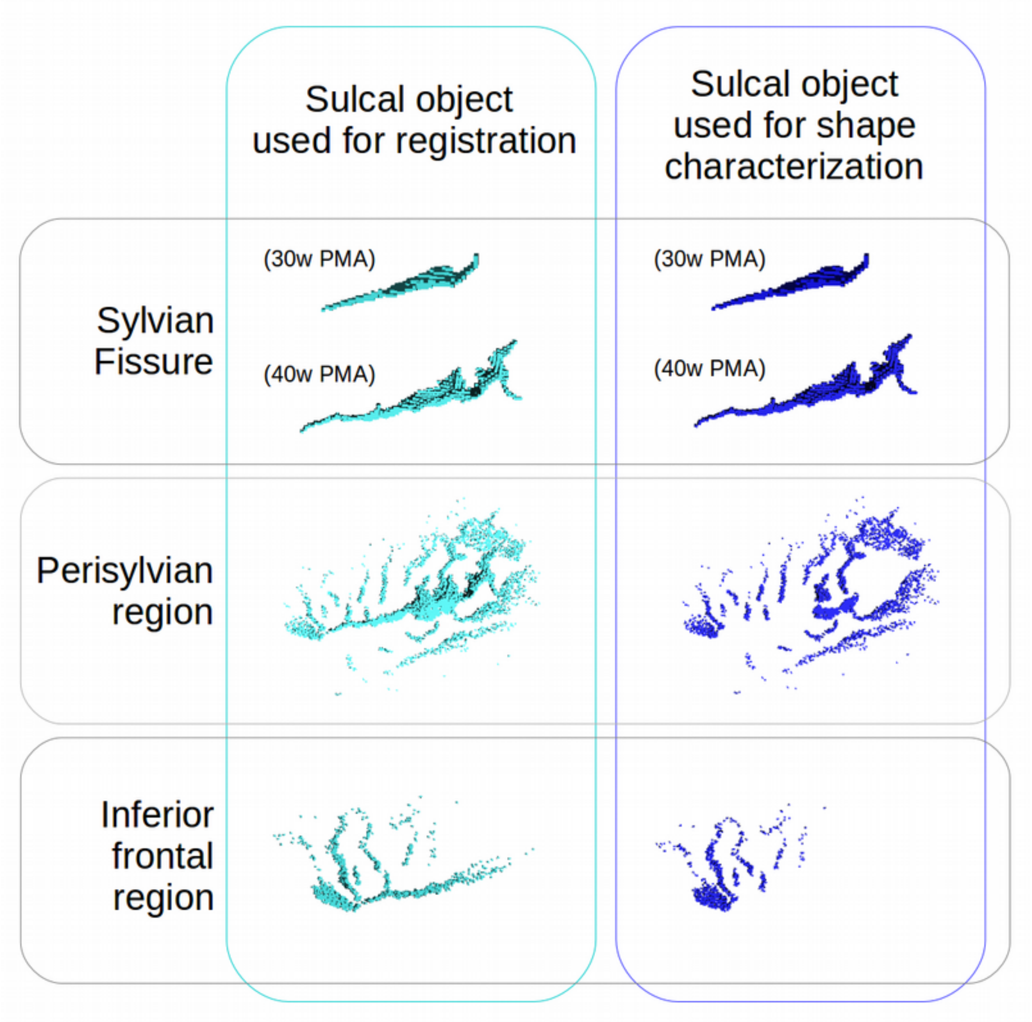


**Supplementary Figure S1. Representation of individual sulcal items used for registration (left panel) and for the analyses of shape characterization (right panel). For the sylvian fissure, one sulcus per age-group was represented and the corresponding age-group was specified. Note that the dark blue point clouds are included in their cyan counterparts.**
